# Smart Leginon enables cryoEM screening automation across multiple grids

**DOI:** 10.1101/2023.07.24.550406

**Authors:** Anjelique Sawh-Gopal, Aygul Ishemgulova, Eugene Y.D. Chua, Mahira F. Aragon, Joshua H. Mendez, Edward T. Eng, Alex J. Noble

## Abstract

**Summary:** CryoEM multi-grid screening is often a tedious process that demands hours of attention. Here, this protocol shows how to set up standard *Leginon* collection and *Smart Leginon Autoscreen* to automate this process. This protocol can be applied to the majority of cryoEM holey foil grids.

Advancements in cryo-electron microscopy (cryoEM) techniques over the past decade have allowed structural biologists to routinely resolve macromolecular protein complexes to near-atomic resolution. The general workflow of the entire cryoEM pipeline involves iterating between sample preparation, cryoEM grid preparation, and sample/grid screening before moving on to high-resolution data collection. Iterating between sample/grid preparation and screening is typically a major bottleneck for researchers, as every iterative experiment must optimize for sample concentration, buffer conditions, grid material, grid hole size, ice thickness, and protein particle behavior in the ice, amongst other variables. Furthermore, once these variables are satisfactorily determined, grids prepared under identical conditions vary widely in whether they are ready for data collection, so additional screening sessions prior to selecting optimal grids for high-resolution data collection are recommended. This sample/grid preparation and screening process often consumes several dozen grids and days of operator time at the microscope. Furthermore, the screening process is limited to operator/microscope availability and microscope accessibility. Here, we demonstrate how to use *Leginon* and *Smart Leginon Autoscreen* to automate the majority of cryoEM grid screening. *Autoscreen* combines machine learning, computer vision algorithms, and microscope-handling algorithms to remove the need for constant manual operator input. *Autoscreen* can autonomously load and image grids with multi-scale imaging using an automated specimen-exchange cassette system, resulting in unattended grid screening for an entire cassette. As a result, operator time for screening 12 grids may be reduced to ∼10 minutes with *Autoscreen* compared to ∼6 hours using previous methods which are hampered by their inability to account for high variability between grids. This protocol and video tutorial first introduces basic *Leginon* setup and functionality, then demonstrates *Autoscreen* functionality step-by-step from the creation of a template session to the end of a 12 grid automated screening session.

## Introduction

Single particle cryo-electron microscopy (cryoEM) allows for near-atomic resolution structure determination of purified macromolecular complexes. A single particle cryoEM experiment only requires one or two well-chosen grids selected from a much larger set of grids with varying sample and grid conditions. Microscope screening to examine these grids entails imaging each grid at several magnifications to determine which grid satisfies most key requirements for a high-resolution data collection, including ice thickness, sufficient areas for full data collection, protein purity, protein concentration, protein stability, and minimal preferred orientation issues^1^. Optimizing for these key requirements often involves feedback between screening at the microscope and preparation conditions such as protein production, buffer selection, potential detergents, and grid type^2–4^ (**Figure 1**). Conventional grid screening is performed manually or semi-manually with software such as Leginon^5^, SerialEM^6^, and EPU^7^. Conventional screening requires the microscope operator to spend hours at the microscope to screen several grids, which creates a significant bottleneck in the high-resolution single particle workflow by occupying the operator with rote operations rather than sample/grid optimization.

**Figure 1:**
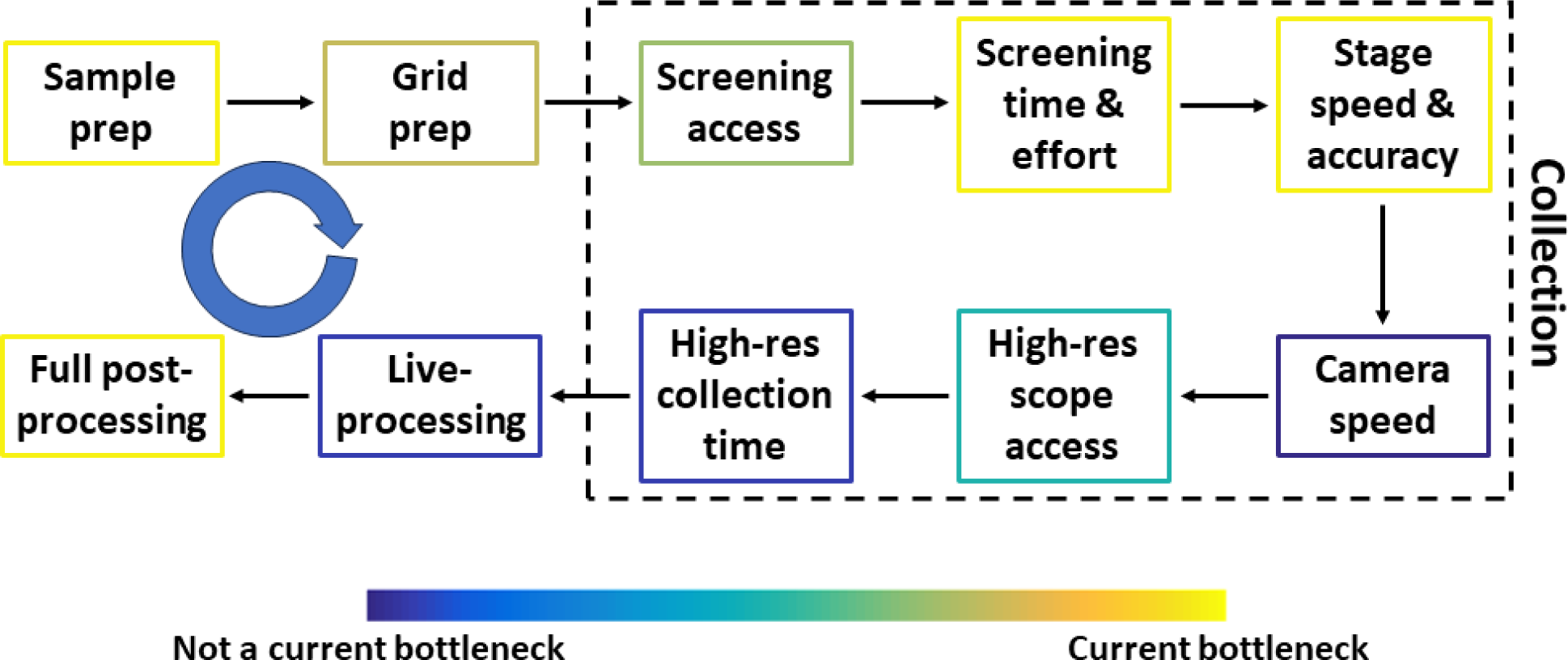
Conventional single particle cryoEM pipeline prior to automated screening. The most common steps in the conventional single particle cryoEM pipeline prior to automated screening, together with components that can be improved. Each step is colored to approximate how much of a bottleneck the step is relative to others. The blue circular arrow represents several feedback loops between most steps. The throughput at several steps depends heavily on the sample, funding, and the researcher’s location.

Previously, *Smart Leginon Autoscreen* and the underlying machine learning software, *Ptolemy*, have been introduced, and their underlying methods and algorithms together with examples have been described^8, 9^. Several other software packages are either capable of or working towards fully automated multi-grid screening^10^, including *SmartScope*^11^, *Smart EPU*^12^, and *CryoRL*^13, 14^. To address the screening bottleneck, *Smart Leginon* allows the user to first set up screening parameters in a template microscope session, then use that template session’s parameters as a template to screen the full cassette of grids in the microscope autoloader. All manual work during the cassette screening is eliminated, which allows for the optimization feedback loop to proceed significantly more efficiently.

In this protocol, the full *Smart Leginon Autoscreen* workflow is described so that the reader may perform fully automated multi-grid cryoEM screening on their own resources. For those new to *Leginon*, the first section in the protocol describes conventional *Leginon* usage. This knowledge is composed of several years of experience across several autoloader microscopes, which is then built upon in the subsequent *Smart Leginon* section of the protocol. Additional tutorial videos may be found on https://youtube.com/playlist?list=PLhiuGaXlZZel9cqXAWsv2lwZmfSQnWqJg

## Protocol

To follow this protocol, depicted in **Figure 2**, *Leginon 3.6+* needs to be installed on the microscope computer and on an additional Linux workstation, and *Ptolemy* needs to be installed on the Linux workstation. This protocol has been developed over several years using Thermo Fisher Scientific (TFS) Glacios and Krios microscopes. This protocol assumes that the reader has already configured *Leginon*, *Appion*^15^, the associated database, microscope calibrations, performed direct alignments on the microscope, and has set up two *Leginon* Applications: One for standard single particle collection and one for single particle collection with *Ptolemy*.

**Figure 2:**
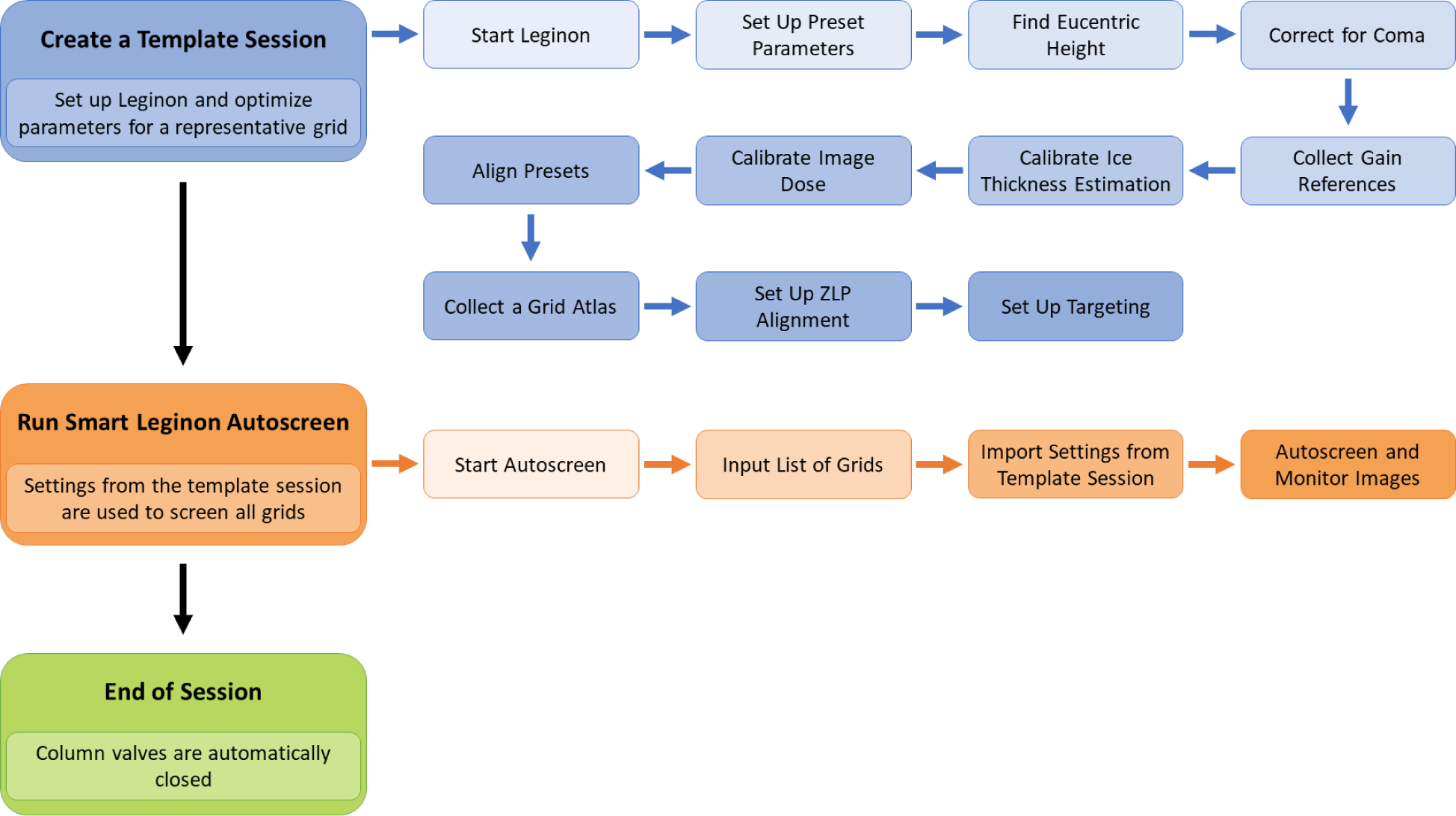
Smart Leginon Autoscreen workflow. A high-level overview of the *Smart Leginon Autoscreen* workflow. First, a template session is created by selecting parameters for a representative grid in the batch of grids to be screened. Setting up *Leginon* and creating a template session takes less than 45 minutes. Second, *Autoscreen* is set up to use the template session parameters to screen all of the grids in the cassette. Setting up *Autoscreen* takes less than 10 minutes. Finally, *Autoscreen* ends the screening session.

Information for setting up *Leginon* is available here: https://emg.nysbc.org/redmine/projects/leginon/wiki/Leginon_Manual. Information for setting up *Ptolemy* within *Leginon* is available here: https://emg.nysbc.org/redmine/projects/leginon/wiki/Multi-grid_autoscreening. Download *Leginon* from http://leginon.org and *Ptolemy* from https://github.com/SMLC-NYSBC/ptolemy.

*Leginon* is licensed under the Apache License, Version 2.0, and Ptolemy is licensed under CC BY-NC 4.0.

### Leginon usage

**1. Start Leginon**
1.1. On the microscope Windows computer, close any *Leginon* clients, then re-open it. In the Linux workstation, open a terminal window and type ‘start-leginon.py’ or the system’s appropriate alias for starting *Leginon*.
1.2. In the new *Leginon* Setup window, select ‘Create a new session’ and click Next.
1.3. Select your project from the dropdown list and click Next.
1.4. Leave the Name as it is and select the correct holder for your microscope setup, then click Next.
1.5. For the Description, enter relevant information such as the microscope name, grid/sample description, and experiment description, then click Next.
1.6. For the Image Directory, make sure the appropriate filesystem is selected, and that the full path is appropriate for saving images, then click Next.
1.7. Under ‘Connect to clients’, click Edit. In the dropdown menu, select all of the computers that should be connected and click the + button for each, then click ‘OK’ and Next.
1.8. Input the correct C2 aperture size and click Next. This value can be found in the ‘Apertures’ tab of TFS TUI software.

**2. Leginon Interface**

2.1. Select Application from the toolbar and click ‘Run…’
2.2. Select the proper Application from the dropdown menu (click ‘Show All’ if necessary). Set ‘main’ to the Leginon computer and ‘scope and camera’ to that respective computer, then click Run.
2.3. The left side of the main *Leginon* window will populate with nodes.

**2.4. Leginon nodes**

2.4.1. The left panel shows all *Leginon* nodes. The green camera icon nodes are the images that will be saved: “Grid”, “Square”, “Hole”, and “Exposure”.
2.4.2. Nodes with the target sign are lower magnification images for targeting the higher magnification images.
2.4.3. The purple camera nodes are the nodes that are programmed to find eucentric z-height and eucentric focus.
2.4.4. Additionally, there are nodes to align the Zero loss peak, monitor buffer cycle, monitor liquid nitrogen filling, collecting gain correction images, calculating ice thickness (IceT), and navigating the grid through different magnifications using stage and image shift.

**2.5. Presets Manager**

2.5.1. Click on the Presets_Manager node. In that node, click the bottom icon for importing presets or the icon above that one to create a new preset from the current state of the microscope.
2.5.2. If the bottom icon is clicked, an ‘Import Presets’ window will open. Select the correct TEM and Digital Camera, then click ‘Find’ and select the most recent session with the desired presets. Highlight all desired presets and click ‘Import’, then click ‘Done’.
2.5.3. The Presets Manager node should now list all imported and created presets. It is recommended to have presets for several magnifications and focusing, including gr: Grid Magnification, sq: Square Magnification, hln: Hole Magnification, fan: Auto-focus, fcn: Central focus, enn: Exposure Magnification (trailing ‘n’ refers to nanoprobe). Typical preset parameters for each magnification are shown in **Tables 1–3**. Note that we use a C2 aperture size of 70 µm for the Glacios, 50 µm for the Krios with a Selectris X and Falcon 4i, and 100 for the Krios with a BioQuantum with a K3.

**Table 1:** Preset parameters for cryoEM grid screening at SEMC using a Glacios with a Falcon 3EC camera. Parameters for each preset commonly used on a Glacios with a Falcon 3EC camera at SEMC are shown. Different microscopes will have varying magnifications available and different experiments will use varying parameters such as defocus and exposure time.

|  | gr: Grid | sq: Square | hln: Hole | fan: Auto-focus | fcn: Central focus | enn: Exposure |
| --- | --- | --- | --- | --- | --- | --- |
| **Magnification** | 210 | 2600 | 6700 | 120000 | 120000 | 120000 |
| **Defocus** | -0.0002 | -0.00015 | -0.00015 | -2e-06 | -7e-07 | -2.5e-06 |
| **Spot size** | 5 | 5 | 4 | 2 | 2 | 2 |
| **Intensity** | 1.1 | 0.83 | 0.65 | 0.44 | 0.44 | 0.45 |
| **Dimension** | 1024 x 1024 | 1024 x 1024 | 1024 x 1024 | 1024 x 1024 | 1024 x 1024 | 4096 x 4096 |
| **Offset** | 0, 0 | 0, 0 | 0, 0 | 0, 0 | 512, 512 | 0, 0 |
| **Binning** | 4 x 4 | 4 x 4 | 4 x 4 | 4 x 4 | 2 x 2 | 1 x 1 |
| **Exposure time (ms)** | 200 | 500 | 500 | 500 | 500 | 1000 |
| **Pre-Exposure (s)** | 0 | 0 | 0 | 0 | 0 | 0 |
| **Dose (e/Å<sup>2</sup>)** | -- | -- | -- | 36.5 | 36.5 | 64.7 |
| **Save raw frames** | No | No | No | No | No | Yes |

**Table 2:** Preset parameters for cryoEM grid screening at SEMC using a Krios with a Selectris X and Falcon 4i camera. Parameters for each preset commonly used on a Krios with a Selectris X energy filter and Falcon 4i camera at SEMC are shown. Different microscopes will have varying magnifications available and different experiments will use varying parameters such as defocus and exposure time.

|  | gr: Grid | sq: Square | hln: Hole | fan: Auto-focus | fcn: Central focus | enn: Exposure |
| --- | --- | --- | --- | --- | --- | --- |
| **Magnification** | 64 | 1700 | 2850 | 75000 | 75000 | 75000 |
| **Defocus** | 0 | -5e-05 | -5e-05 | -1e-06 | -7e-07 | -2e-06 |
| **Spot size** | 6 | 9 | 9 | 6 | 6 | 7 |
| **Intensity** | 0.001 | 1.65e-05 | 1.5e-05 | 4.3e-07 | 4.3e-07 | 5.5e-07 |
| **Energy filter width** | -- | -- | -- | 20 | 20 | 20 |
| **Dimension** | 1024 x 1024 | 1024 x 1024 | 1024 x 1024 | 1024 x 1024 | 2048 x 2048 | 4096 x 4096 |
| **Offset** | 0, 0 | 0, 0 | 0, 0 | 0, 0 | 0, 0 | 0, 0 |
| **Binning** | 4 x 4 | 4 x 4 | 4 x 4 | 4 x 4 | 2 x 2 | 1 x 1 |
| **Exposure time (ms)** | 500 | 2000 | 1000 | 500 | 300 | 8700 |
| **Pre-Exposure (s)** | 0 | 0 | 0 | 0 | 0 | 0 |
| **Dose (e/Å<sup>2</sup>)** | -- | -- | -- | -- | -- | 47.4 |
| **Save raw frames** | No | No | No | No | No | Yes |

**Table 3:** Preset parameters for cryoEM grid screening at SEMC using a Krios with a BioQuantum and K3 camera. Parameters for each preset commonly used on a Krios with a BioQuantum energy filter and K3 camera at SEMC are shown. Different microscopes will have varying magnifications available and different experiments will use varying parameters such as defocus and exposure time.

|  | gr: Grid | sq: Square | hln: Hole | fan: Auto-focus | fcn: Central focus | enn: Exposure |
| --- | --- | --- | --- | --- | --- | --- |
| **Magnification** | 1550 | 940 | 2250 | 81000 | 81000 | 81000 |
| **Defocus** | 0 | -5e-05 | -5e-05 | -1e-06 | 7e-07 | -2e-06 |
| **Spot size** | 4 | 8 | 7 | 6 | 6 | 6 |
| **Intensity** | 0.0015 | 0.00017 | 7.3e-05 | 1.3e-06 | 1.3e-06 | 9.2e-07 |
| **Energy filter width** | -- | -- | 50 | 20 | 20 | 20 |
| **Dimension** | 1024 x 1024 | 1440 x 1024 | 1440 x 1024 | 1440 x 1024 | 1008 x 1008 | 5760 x 4092 |
| **Offset** | 0, 0 | 0, 0 | 0, 0 | 0, 0 | 936, 519 | 0, 0 |
| **Binning** | 4 x 4 | 8 x 8 | 8 x 8 | 8 x 8 | 4 x 4 | 2 x 2 |
| **Exposure time (ms)** | 250 | 600 | 600 | 500 | 500 | 2100 |
| **Pre-Exposure (s)** | 0 | 0 | 0 | 0 | 0 | 0 |
| **Dose (e/Å<sup>2</sup>)** | -- | -- | -- | -- | -- | 51 |
| **Save raw frames** | No | No | No | No | No | Yes |

**2.6. Navigation and eucentric height**

2.6.1. To become familiar with controlling the microscope through *Leginon* and to set the z-height of the grid, go to the Navigation node, select the ‘gr’ preset at the top, and click on the red arrow to the right to send the preset settings to the microscope. The microscope should update after one or two seconds. Once updated, click the camera button to the right to acquire an image.
2.6.2. Using the cursor tool, select a grid square for where to move the stage. Send the sq magnification to the microscope and acquire an image.
2.6.3.Go to the Z_Focus node and click the ‘Simulate Target’ button at the top near the middle of the buttons. While images are being collected for stage tilt focusing, switch to correlation view and watch the peak to ensure that it is in a corner of the correlation image. Once focusing is finished, the stage should be set to the grid’s z-height.

**2.7. Coma correction**

2.7.1. This sub-section assumes that direct alignments have already been performed and that coma corrections have not been performed outside of *Leginon*.
2.7.2. Navigate to an area of the grid that produces clear Thon rings, such as carbon substrate. Note: A cross-grating may be used if collection is to be done on a gold grid.
2.7.3. In the Beam_Tilt_Image settings, the ‘Presets Order’ should include only ‘fcn’.
2.7.4. Click ‘Simulate Target’ to create a Zemlin tableau. Click ‘Tableau’ on the left side of the main window to view the tableau.
2.7.5. Correct for coma by comparing the left and right Fourier transforms with each other and the top and bottom Fourier transforms with each other. If the pairs of images are not identical, first click the cursor icon to the right of the image adjustments, then click slightly off of center in the Tableau image in the direction of the difference and wait for a new set of Fourier transforms to collect.

**2.8. Gain references**

2.8.1. In the Navigation node, send a low magnification preset, such as ‘gr’, and navigate to an area where there is vacuum.
2.8.2. Confirm that the stage position is under vacuum by taking a medium magnification image using the ‘sq’ or ‘hln’ preset.
2.8.3. Send the high magnification ‘enn’ preset to the microscope.
2.8.4. In the Correction node settings, select the proper ‘Instrument’ information and set up the ‘Camera Configuration’ to match your collection settings.
2.8.5. Collect a dark reference image by closing the column valves on the microscope, then in the Correction node select ‘Dark’ and ‘Both Channels’ from the dropdown menus at the top, and click the ‘Acquire’ camera button to the right.
2.8.6. Once complete, select ‘Bright’ from the dropdown menu and click ‘Acquire’. *Leginon* will open the column valves automatically.
2.8.7. Check that the gain was properly collected by selecting ‘Corrected’ from the dropdown menu, clicking ‘Acquire’, and observing the resulting image.

**2.9. Ice thickness reference image**

2.9.1. If the microscope has an energy filter, then in the IceT node settings, check ‘Collect ice thickness image’, enter 395 for the ‘Mean free path’ and fill in the rest of the values for your setup.
2.9.2. If the microscope does not have an energy filter, then in the Navigation node, send the ‘enn’ preset to the microscope and click ‘Acquire’. Make note of the ‘Mean’ pixel value on the left side. in the IceT node settings, check ‘Calculate ice thickness from aperture limited scattering’, enter 1055 for the ‘ALS coefficient’ and the measured mean pixel value.

**2.10. Image dose calibration**

2.10.1. In the Preset_Manager, select the ‘enn’ preset and click the camera button ‘Acquire dose image for the selected preset’. Check the measured dose on the bottom. If it is close to the expected value (typically between 30 and 70), click ‘YES’.

**2.11. Preset alignments**

2.11.1. In the Preset_Manager, check all of the high magnification presets (‘enn’, ‘fcn’, and ‘fan’) to make sure the image shift and beam shift are 0, 0.
2.11.2. On the microscope computer, navigate to a carbon area.
2.11.3. On the *Leginon* computer in the Navigation node, acquire an image with the ‘gr’ preset.
2.11.4. Find an object of interest and go to that location using the cursor tool.
2.11.5. Acquire an image with the ‘hln’ preset and relocate a unique part of that object of interest to the center using the cursor tool.
2.11.6. Acquire an image with the ‘enn’ preset and relocate to the same unique part of the object of interest using the cursor tool.
2.11.7. Select ‘image shift’ from the pulldown menu and acquire an image with the ‘hln’ preset. Relocate to the same unique part of the object of interest with the cursor tool.
2.11.8. In the Presets_Manager, select the “hln” preset, click on the settings button, and import the image shift from Navigation by clicking the left green arrow next to the ‘Image shift’ values.
2.11.9. Repeat steps 2.11.7 and 2.11.8 for the ‘sq’ and ‘gr’ presets.

**2.12. Grid atlas**

2.12.1. On the microscope computer, close column valves and retract the objective aperture. Go to the Grid_Targeting node. In settings, change the label of the grid. Choose the desired radius of your atlas (the maximum radius is 0.0009 m). Click ‘OK’. Then click the ‘Calculate atlas’ calculator button at the top and click the green Play button (‘Submit Targets’).
2.12.2. In the Square_Targeting node, the grid images will be collected and stitched together to form an atlas. You can zoom in and out using the dropdown menu and adjust contrast and brightness. You can use the scroll bars to move throughout the grid.
2.12.3. Once the atlas is collected, insert the objective aperture if desired.
2.12.4. If the microscope has an energy filter, select a reference target in the center of a broken square, press the ‘Play’ button, and proceed with ZLP alignment in the following sub-section, otherwise skip the ZLP alignment step.

**2.13. ZLP alignment**

1. 2.13.1. In the Align_ZLP node settings, select ‘stage position’ to move the reference target and select ‘preset manager’ as the mover. Deselect ‘bypass conditioner’, then press OK.
2. 2.13.2. ZLP alignment should now be configured so that the microscope periodically moves to the reference target and executes the camera’s ZLP alignment routine.

**2.14. Hole template targeting setup**

2.14.1. In the Square_Targeting node, select multiple acquisition targets, then press Play.
2.14.2. In the Hole_Targeting node settings, make sure ‘Allow user verification of selected targets’ and ‘Queue up targets’ are checked. Also, check ‘Skip automated hole finder’ for now. Click Apply, then OK.
2.14.3. In the main window, use Ctrl-Shift-right click to remove all targets. Select the acquisition cursor and place your own targets. Select the focus cursor and place a focus target in between the acquisition targets. Click ‘Play’.
2.14.4. For the next Hole_Targeting image, uncheck the ‘Skip automated hole finder’ in settings, then click ‘Apply’ and ‘OK’. Remove the auto targets with Ctrl-Shift-right click.
2.14.5. Select the ruler tool and measure the diameter across a hole. In Template settings, change the ‘Final Template Diameter’ to the measured hole diameter. Do not change the ‘Original Template
2.14.6. Diameter’. Click ‘Test’. If bright peaks are not in the center of each hole, increase the ‘Final Template Diameter’. When finished, click ‘OK’.
2.14.7. In the Threshold settings, choose a value for ‘A’ that segments the holes individually when ‘Test’ is clicked, as shown in the video. Click ‘OK’ when satisfied.
2.14.8. In the Blobs settings, input the values shown in the video and click ‘Test’. The Max blobs value is 1 so only one blob shows up. Click ‘OK’.
2.14.9. In Lattice settings, use the ruler tool to measure the distance between two holes and input the value in ‘Spacing’ and click ‘Test’. The one blob will turn into a lattice point. Click ‘OK’.
2.14.10. Go to acquisition settings, optimize the acquisition targets using ice thickness thresholds and the ‘Test targeting’ button. Get ice thickness information by hovering over the lattice points.
2.14.11. If the acquisition targets aren’t satisfactory, use the ruler tool to measure the distance and angle from the lattice point to the desired location for an acquisition target. Delete the previous ‘Acquisition Target Template’ points. Click on ‘Auto Fill…’, put 4 for the number of targets, and change the radius and angle to the measured values. Click ‘OK’. Check ‘Apply ice thickness threshold on template-convolved acquisition targets’.
2.14.12. Once satisfied with the lattice points and the ice thickness thresholds, click the ‘Submit Targets’ button.
2.14.13. Repeat any of the above steps for each square that was selected. Submit the whole queue with the ‘Submit Queued Targets’ button once all the square targets are submitted.
2.14.14. *Leginon* will begin focusing and imaging each set of targets. In the Z_Focus node, make sure that the eucentric height is found properly.

**2.15. Exposure template targeting setup**

2.15.1. In the Exposure targeting node, Hole Magnification images will appear. Use Ctrl-Shift-right click to remove the auto targets.
2.15.2. Measure the diameter of a hole with the ruler tool. In the Template settings, input the diameter into ‘Final Template Diameter’ and click ‘Test’. A peak should now be at the center of each hole. Adjust the ‘Diameter’ values if necessary.
2.15.3. In the Threshold settings, adjust the ‘A’ value until the binarized Test image shows white areas only where holes are located.
2.15.4. In the Blobs settings, click ‘Test’. One blob per segmented hole should appear. Increase the ‘Border’ to remove the blobs from the edges of the image, if desired.
2.15.5. In the Lattice settings, click ‘Test’. Adjust parameters until all blobs have turned into lattice points. Click ‘OK’.
2.15.6. Click on the ruler tool and measure the distance between two lattice points. In the Lattice settings, change ‘Spacing’ to that distance.
2.15.7. Hover over each lattice point to see the mean intensity, mean thickness, standard deviation intensity, and standard deviation thickness. Make note of the intensities for each lattice point and use them to set desired ice thickness parameters in the acquisition settings.
2.15.8. Measure distance and angle from one lattice point to the center of 4 holes with the ruler tool. In the acquisition settings, delete the current focus targets. Click ‘Auto Fill…’, change the radius and angle to the measured values. Click ‘Test targeting’, click ‘OK’, and click ‘Submit Targets’.
2.15.9. *Leginon* will find eucentric focus (Focus node) and collect exposures, which will appear in the Exposure node.
2.15.10. Once all targets are imaged, go to the Exposure_Targeting node to see the next hole image. In the settings, uncheck ‘Allow for user verification of selected targets’. Click ‘OK’ and click ‘Submit Targets’.
2.15.11. *Leginon* will automatically collect images based on the settings configured above. Images and metadata can be seen in *Appion*.
2.15.12. Changes can be made during automated collection. For example, the collection defocus range can be changed at any time by editing the ‘enn’ preset in the Preset_Manager.
2.15.13. If collection needs to be stopped, the queue can be ended by clicking the ‘Abort’ and ‘Abort Queue’ buttons in the Hole and Exposure nodes.
2.15.14. Once *Leginon* is done collecting, go to Application and click Kill, then go to File and click Exit.

### Smart Leginon Autoscreen usage

**3. Create a Smart Leginon Template Session**

3.1. Follow the instructions in Section 1 to start *Leginon*.
3.2. Go to Application and click ‘Run…’. In the Run Application window, select the Ptolemy Application (select ‘Show All’ if necessary). Set ‘main’ to the Leginon computer and ‘scope and camera’ to that respective computer.
3.3. In the Preset_Manager, import presets as described in Section 2.5.

**3.4. Configure node settings**

3.4.1. In the Square_Targeting node settings, make sure ‘Sort targets by shortest path’ and ‘Enable auto targeting’ are checked (**Supplementary** Figure 1a).
3.4.2. In the Square node settings, make sure ‘Wait for node to process the image’ is checked. Add the Square preset to the list on the right from the dropdown menu if it is not already there. In Advanced settings, check ‘set these apertures while imaging’ and ensure that the values for the two apertures are correct (**Supplementary** Figure 1b).
3.4.3. In the Hole_Targeting node settings, check ‘Allow for user verification of selected targets’. ‘Queue up targets’ and ‘Skip automated hole finder’ should be unchecked (**Supplementary** Figure 2a).
3.4.4. In the Hole node setting, check ‘Wait for a node to process the image’ and the Hole preset is in the list on the right. In Advanced settings, check ‘set these apertures while imaging’ and ensure that the values for the two apertures are correct (**Supplementary** Figure 2b).
3.4.5. In the Exposure_Targeting node settings, check ‘Allow for user verification of selected targets’. ‘Queue up targets’ and ‘Skip automated hole finder’ should be unchecked (**Supplementary** Figure 3a).
3.4.6. In the Exposure node settings, make sure ‘Wait for a node to process the image’ is unchecked, the Exposure preset is listed on the right, and in Advance settings check ‘set these apertures while imaging’ and ensure that the values for the two apertures are correct (**Supplementary** Figure 3b).
3.4.7. In the Focus node settings, make sure ‘Wait for a node to process the image’ is unchecked, the Auto-focusing preset is listed on the right, and the ‘Desired autofocus accuracy’ is set to 4e-6 m (**Supplementary** Figure 4a).
3.4.8. In the Focus node Focus Sequence (next to the settings button), enable only two Beam Tilt Auto-focusing steps (**Supplementary** Figure 4b,c).
3.4.9. In the Z_Focus node settings, make sure ‘Wait for a node to process the image’ is unchecked, the Hole preset is listed on the right, and ‘Desired autofocus accuracy’ is 5e-5 m (**Supplementary** Figure 4a).
3.4.10. In the Z_Focus node Focus Sequence, enable only two low-magnification Stage Tilt steps (**Supplementary** Figure 5b,c).
3.4.11. Determine the grid’s z-height as described in Section 2.6.
3.4.12. Collect an atlas as described in Section 2.7.

**3.7. Set up square finder parameters**

3.7.1. Once the atlas has been collected, *Ptolemy* will locate squares in the Square_Targeting node. Each square will show a blue circle, called a blob. When hovering over each blob, *Leginon* will report their size as calculated by *Ptolemy*. Make note of the largest and smallest blobs.
3.7.2. In the Thresholded settings, change the minimum and maximum ‘Filter Range’ to include desirable squares and exclude undesirable squares.
3.7.3. Click the Find Squares button in the top toolbar. Adjust the Filter Range until Find Squares targets well.
3.7.4. In the acquisition settings, choose values for the ‘Max. number of targets’ and ‘Number of target group to sample’. These parameters will define how many squares and groups of squares are targeted.
3.7.5. Once satisfied with the parameters, click the Play button. An example atlas after setup is shown in **Supplementary** Figure 6.

**3.8. Set up hole finder parameters**

3.8.1. In the Hole_Targeting node, use the ruler tool to measure the diameter of a hole.
3.8.2. In the Template settings, input the diameter into ‘Final Template Diameter’ and click ‘Test’. Adjust the diameter until all holes have bright white peaks in the center.
3.8.3. In the Threshold settings, click ‘Test’. Adjust the ‘A’ value until the binarized image shows white areas only where holes are located.
3.8.4. In the Blobs settings, ‘Border’ targets may be excluded by using the ruler to determine a minimum distance from the edge and inputting that value. Blobs may be filtered by their size, roundness, and number desired. Hover over blobs to show their values. Click ‘Test’ to inspect current values.
3.8.5. In the Lattice settings, input the radius of the holes and spacing between holes (use the measuring tool), then click the ‘42’ button to measure the ‘Reference Intensity’ value of a vacuum area (empty hole or broken support film).
3.8.6. In the acquisition settings, check ‘Use subset of the acquisition targets’ and set the ‘Sample Maximal’ value to a small number, such as 2. Set a wide range of ice thickness means and standard deviations (measure values by hovering over targets). Click ‘Test targeting’ to randomize the target selection given the values above.
3.8.7. Click the Play button when satisfied with all settings. *Leginon* will perform stage Z_Focus and collect the first target. An example image after setup is shown in **Supplementary** Figure 7.

**3.9. Set up exposure targeting parameters**

3.9.1. In the Hole settings, set the Shell Script to the hl_finding.sh script path in the *Ptolemy* installation. Set the minimum score to accept to be ≤0. Input the radius of the holes (use measuring tool), then click the ‘42’ button to measure the ‘Reference Intensity’ value of a vacuum area (empty hole or broken support film). Click ‘Test’ to find the lattice of holes.
3.9.2. In acquisition settings, check ‘Use subset of the acquisition targets’ and set the ‘Sample Maximal’ value to a small number, such as 4, to collect on a subset of holes for screening. Set a wide range of ice thickness means and standard deviations (measure these values by hovering over targets).
3.9.3. Click the Play button when satisfied with all settings. *Leginon* will perform eucentric Focus and collect high-magnification images, which can be seen in the Exposure node. An example image after setup is shown in **Supplementary** Figure 8.
3.9.4. Check the next Exposure_Targeting image to see if the settings above are still sufficient. Once satisfied, uncheck ‘Allow for user verification of selected targets’ in the Exposure Targeting and Hole Targeting settings.
3.9.5. Screening should now be running unattended for the current grid. This session will be used as the template session for all grids.
4. Once the grid is done screening, click File > Exit to close *Leginon*.

**5. Set up Smart Leginon Autoscreen**

5.1. In a terminal window, execute *Smart Leginon* autoscreen.py.
5.2. Select ‘gui’, input a comma-separated list of grid slots to screen, select full workflow, enter the template session name to base new sessions on (this can be found in *Appion* imageviewer), and use the template session’s z-height value.
5.3. A gui will open to allow you to enter the session name for each grid and select their respective project associations.
5.4. *Smart Leginon Autoscreen* will now use the template session settings to automatically screen each grid and switch between grids unattended.
5.5. The user may follow along during the collection in *Leginon*, *Appion*, and the microscope computer, or may leave the microscope completely unattended.
6. Once all grids are screened, *Smart Leginon* will close the column valves on the microscope.

### Representative Results

Following the protocol, cryoEM screening sessions may be run automatically and successfully for the majority (80-90%) of holey grids and conditions. Several examples and experiments have been presented previously^8, 9^ to demonstrate the expected outcomes of successful *Smart Leginon Autoscreen* sessions. A successful *Autoscreen* session begins with ∼10 minutes of setup and commonly results in a full cassette of grids screened automatically after about 6 hours (30 minutes per grid) where 3-5 squares of different size and 3-5 holes per square are imaged at high magnification, allowing the user to quickly determine the characteristics of the sample on each grid and rapidly iterate through sample/grid conditions (**Figure 3**). Occasionally sessions are unsuccessful, commonly due to *Autoscreen* targeting broken squares, not interpreting large ice thickness gradients across the grid or across squares properly, or failing to identify holes properly on carbon grids. Additionally, potential memory leaks may cause *Leginon* to crash due to excessive memory usage, which may be solved by freeing up RAM or rebooting the computer, or ameliorated by adding more RAM to the computer.

**Figure 3:**
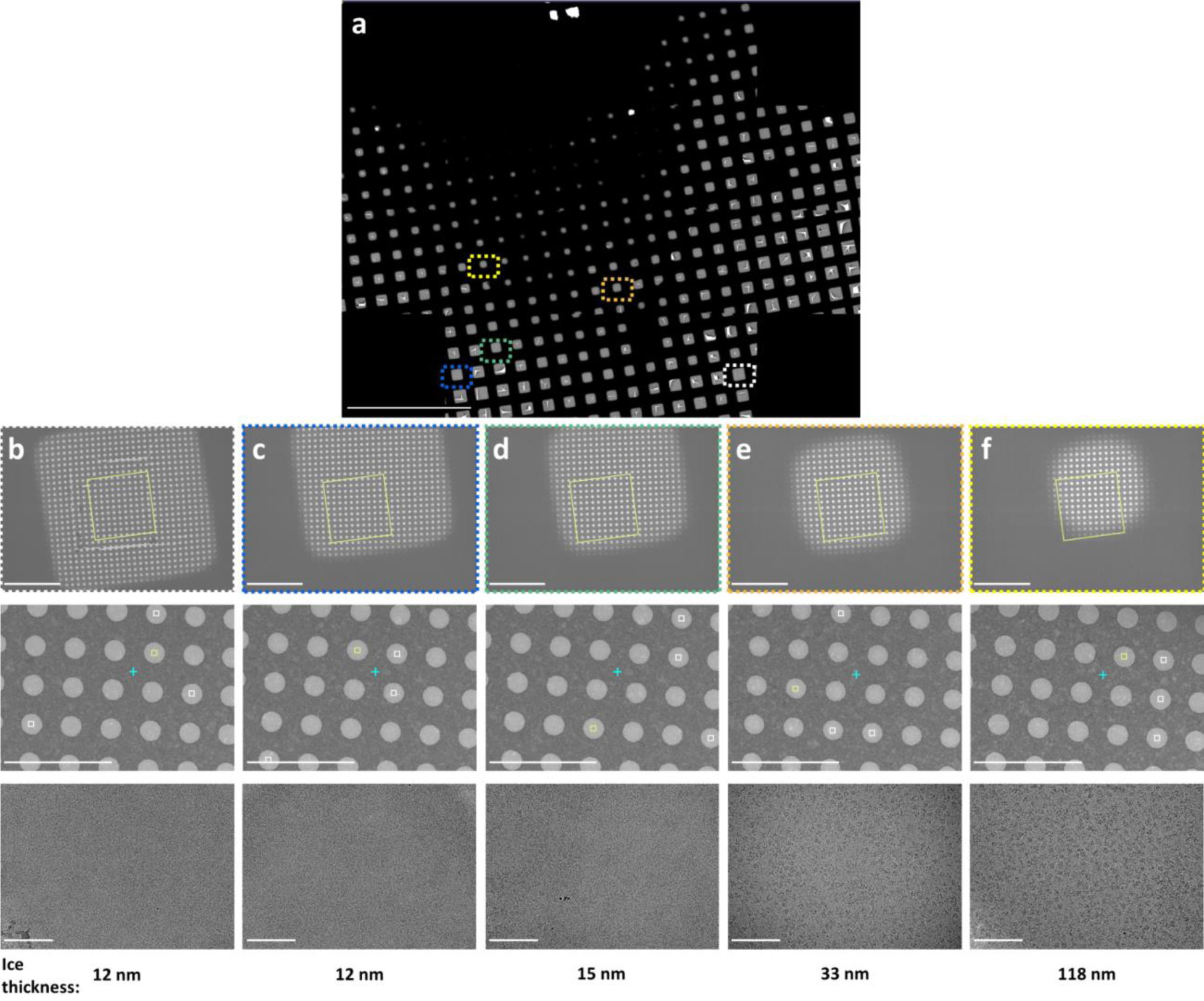
Representative Smart Leginon Autoscreen results. Representative multi-scale images following the *Smart Leginon Autoscreen* protocol collected on a TFS Krios cryoTEM with a BioQuantum energy filter and K3 camera. (a) A composite ‘atlas’ image showing an overview of a cryoEM grid. (b)-(f) Multi-scale images from indicated locations in the grid atlas. Low magnification images in the first row, medium magnification images in the second row, and high magnification images in the third row were each selected automatically to obtain information about the sample from thin to thick ice squares. Ice thickness as estimated by *Leginon* is shown on the bottom. Scale bars are 500 µm in (a) and 10 µm for the first row, 5 µm for the second row and 100 nm for the third row for (b)-(f). Modified from Cheng et al., 2023^8^ under CC BY 4.0.

## Discussion

In this protocol, we describe the pipeline for *Smart Leginon Autoscreen*, and additionally basic *Leginon* usage for those new to the collection software. Single particle cryoEM is poised to become the most productive 3D protein structure resolving technique by the end of 2024^16^. The single particle cryoEM pipeline consists of several steps that are constantly being optimized to increase data quality and throughput. **Figure 2** shows the most common steps (sample preparation, grid preparation, screening time and effort, high-resolution collection time, live processing, and full post-processing) along with other components of the pipeline that can be improved (screening microscope access, stage speed and accuracy, camera speed, and high-resolution microscope access). Results from most steps become feedback loops into previous steps (blue arrows in **Figure 2**), making the entire pipeline highly interdependent. Each step in **Figure 2** is colored to approximate how much of a bottleneck the step is relative to others. *Smart Leginon Autoscreen* significantly reduces operator time and effort for screening 12 grids from 6 hours to less than 10 minutes, thus relieving that bottleneck and allowing for more rapid feedback to sample/grid preparation (**Figure 3**).

There are several critical steps in the Protocol, depicted in **Figure 1**. It is critical that the grid used for creating the template session is representative of the remaining grids to be screened. Importantly, *Leginon* remembers all settings in the entire setup process for creating a template session (blue steps in **Figure 1**), which allows for recurrent template sessions to be set up more quickly each time. When creating a template session, the most critical step is setting up targeting at all magnifications so that the parameters and thresholds reflect the expected variation across the grids to be screened. The various ‘Test’ buttons allow for efficiency in this setup process. During an *Autoscreen* session, it is critical to monitor the first few grids in *Appion* to quickly detect any issues and fix them inside of *Leginon* as soon as possible.

The typical workflow at SEMC is to feed *Autoscreen* data into CryoSPARC Live^17^ and use this additional information to inform the feedback loops into sample/grid preparation. During intensive researcher-operator cryoEM optimization days, information about the sample and grid conditions is fed back into sample and grid preparation while *Autoscreen* is still screening grids. This allows for several dozen grids to be frozen and screened per week^8^.

*Smart Leginon Autoscreen* works for the majority (80-90%) of holey grids and conditions observed at SEMC. The remaining 10-20% of grids include those that sometimes do not work well – grids with minimal contrast difference between holes and substrate; grids with smaller holes and spacing (e.g., 0.6/0.8) – and grids where targeting across multiple grids is often impractical – Spotiton/Chameleon^18, 19^ grids that consist of stripes of sample across the grid; lacey grids. It may be possible to modify the protocol to work with Spotiton/Chameleon grids by first imaging areas of the stripe manually to determine narrow parameter thresholds, then attempting to group larger and smaller squares together, respectively, in **Step 3.7.4**, and then selecting targets from the group with ice. The goal of this modification is to have *Smart Leginon* separate empty and non-empty squares into two groups. If parameters are found, they may not extend well to the remaining grids to be screened. It may also be possible to modify the protocol to work with lacey grids by removing the hl_finding.sh script in **Step 3.9.1** and configuring the parameters to target lighter/darker areas as desired. The success rate of this modification may vary from grid to grid based on ice thicknesses and grid material.

Troubleshooting during an *Autoscreen* session is possible and sometimes appropriate. Changes to preset (e.g., defocus) and targeting parameters (e.g., Hole Targeting thresholds) can be made during automated collection. While an *Autoscreen* session is collecting, a grid session cannot be canceled because it will terminate autoscreen.py. However, the Abort buttons in the Targeting nodes may be used to skip any part of a grid or an entire grid. Occasionally, autoscreen.py may use too much memory and freeze, offering two options: ‘force quit’ or ‘wait’. If ‘force quit’ is selected, the entire script will terminate, requiring the user to rerun the script to be applied to the remaining grids for screening. If ‘wait’ is selected, the script will continue, and settings may be altered to prevent future freezing, e.g., turning off the image display in the Exposure node, decreasing pixel size in the atlas, or running a memory clear script. If the program freezes without offering the two options, memory errors may not resolve on their own, causing a pause in acquisition. The ‘force quit’ option may be useful in this instance.

*Smart Leginon Autoscreen* is used regularly at SEMC. As bottlenecks in the single particle cryoEM pipeline continue to be reduced, cryoEM adoption will continue to increase to help answer biological questions. This Protocol is a step in the direction of optimizing the entire pipeline by providing a clear path for significantly reducing feedback loops.

## Acknowledgements

Some of this work was performed at the Simons Electron Microscopy Center at the New York Structural Biology Center, with support from the Simons Foundation (SF349247), NIH (U24 GM129539), and NY State Assembly.

---

**Supplementary Figure 1:**
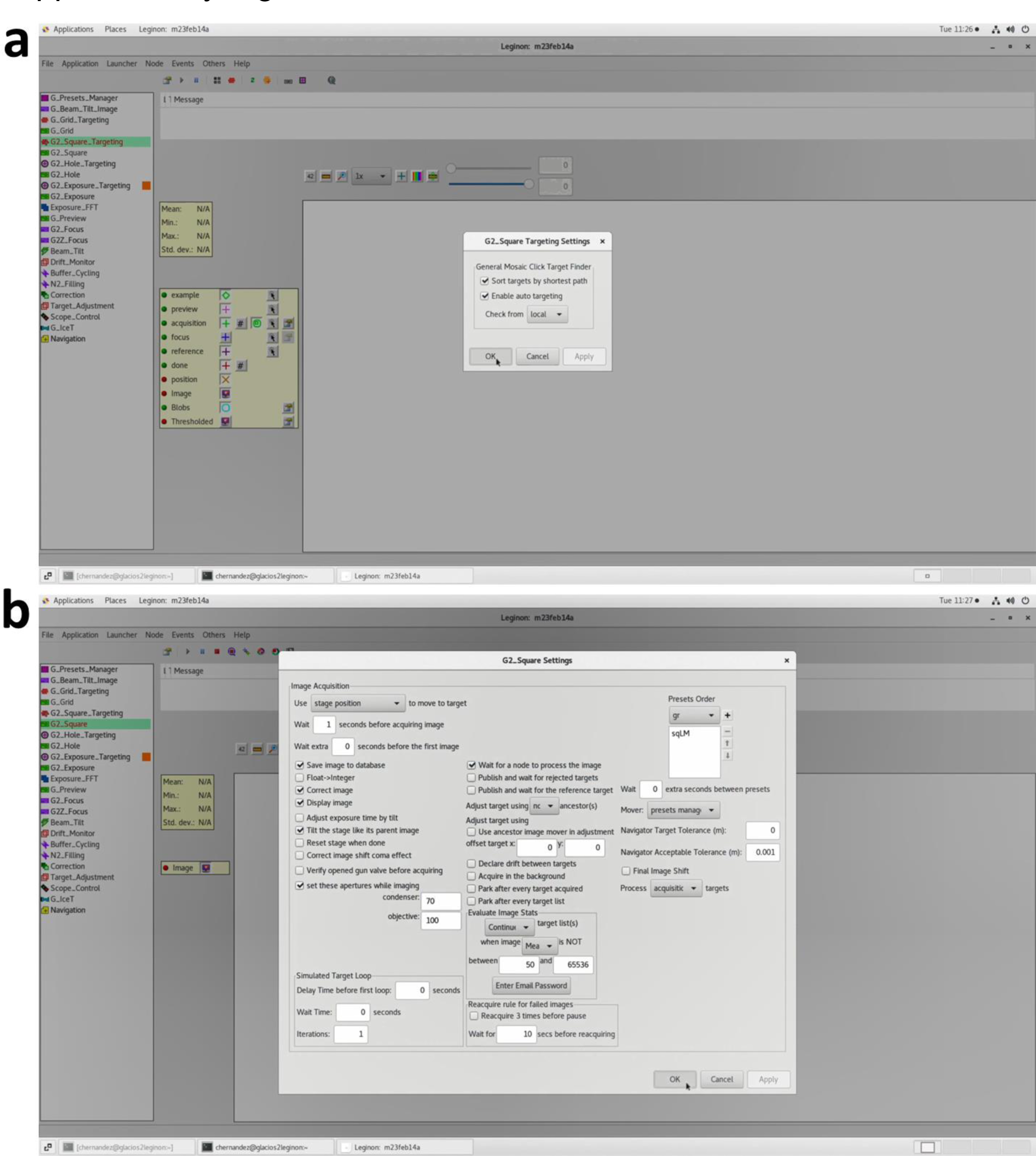
Square Targeting settings and Square settings for *Smart Leginon*.

**Supplementary Figure 2:**
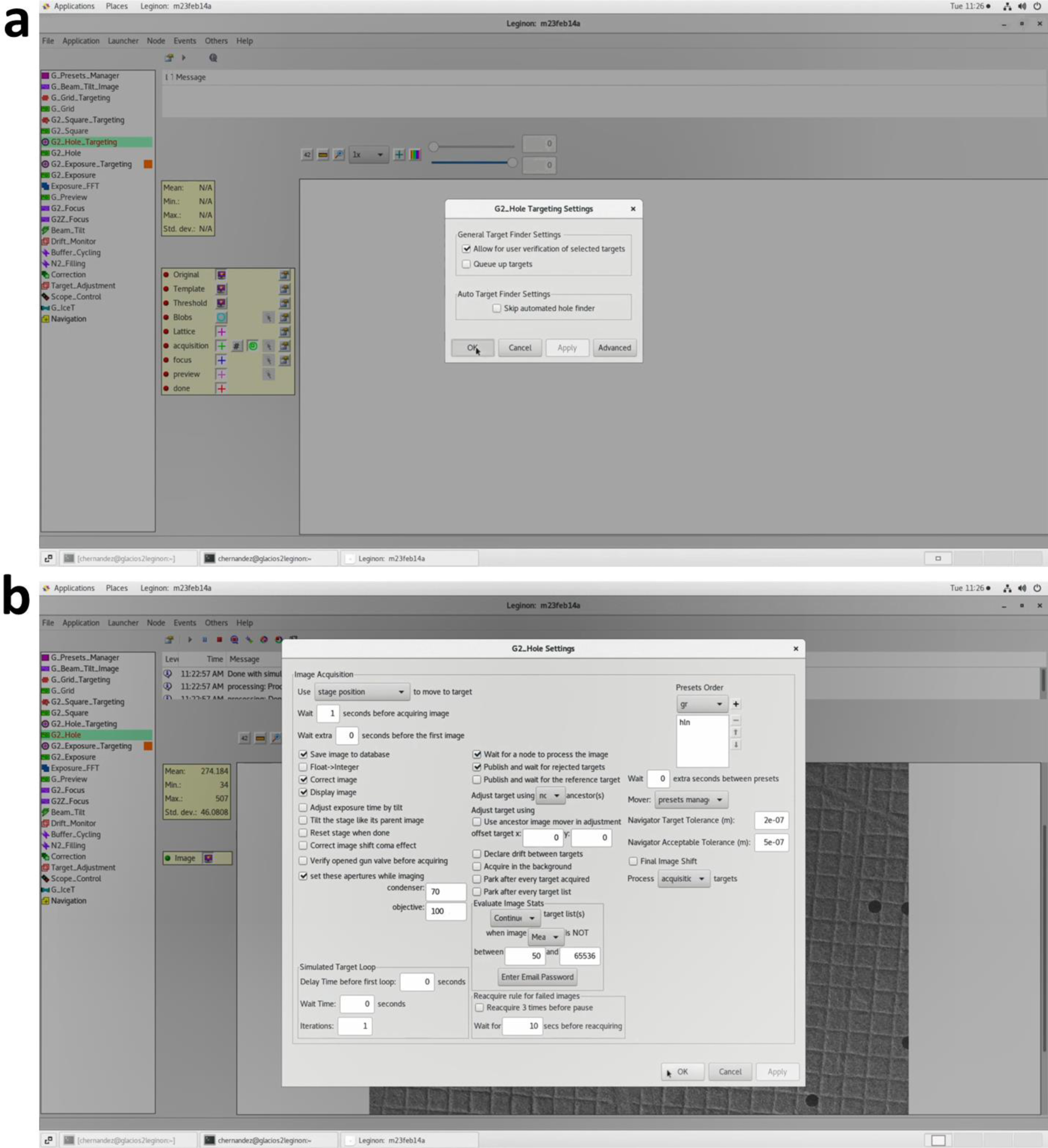
Hole Targeting settings and Hole settings for *Smart Leginon*.

**Supplementary Figure 3:**
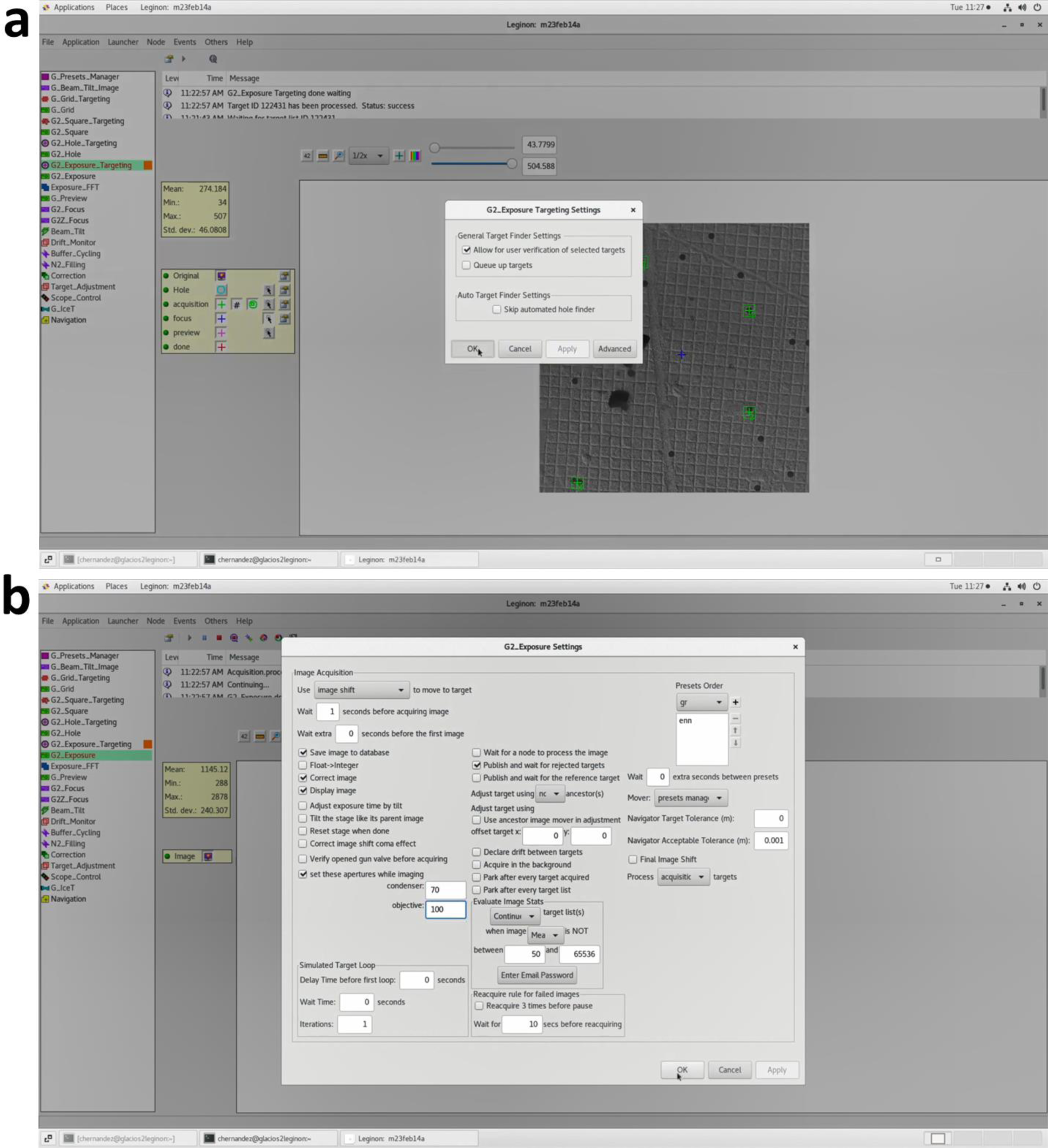
Exposure Targeting settings and Exposure settings for *Smart Leginon*.

**Supplementary Figure 4:**
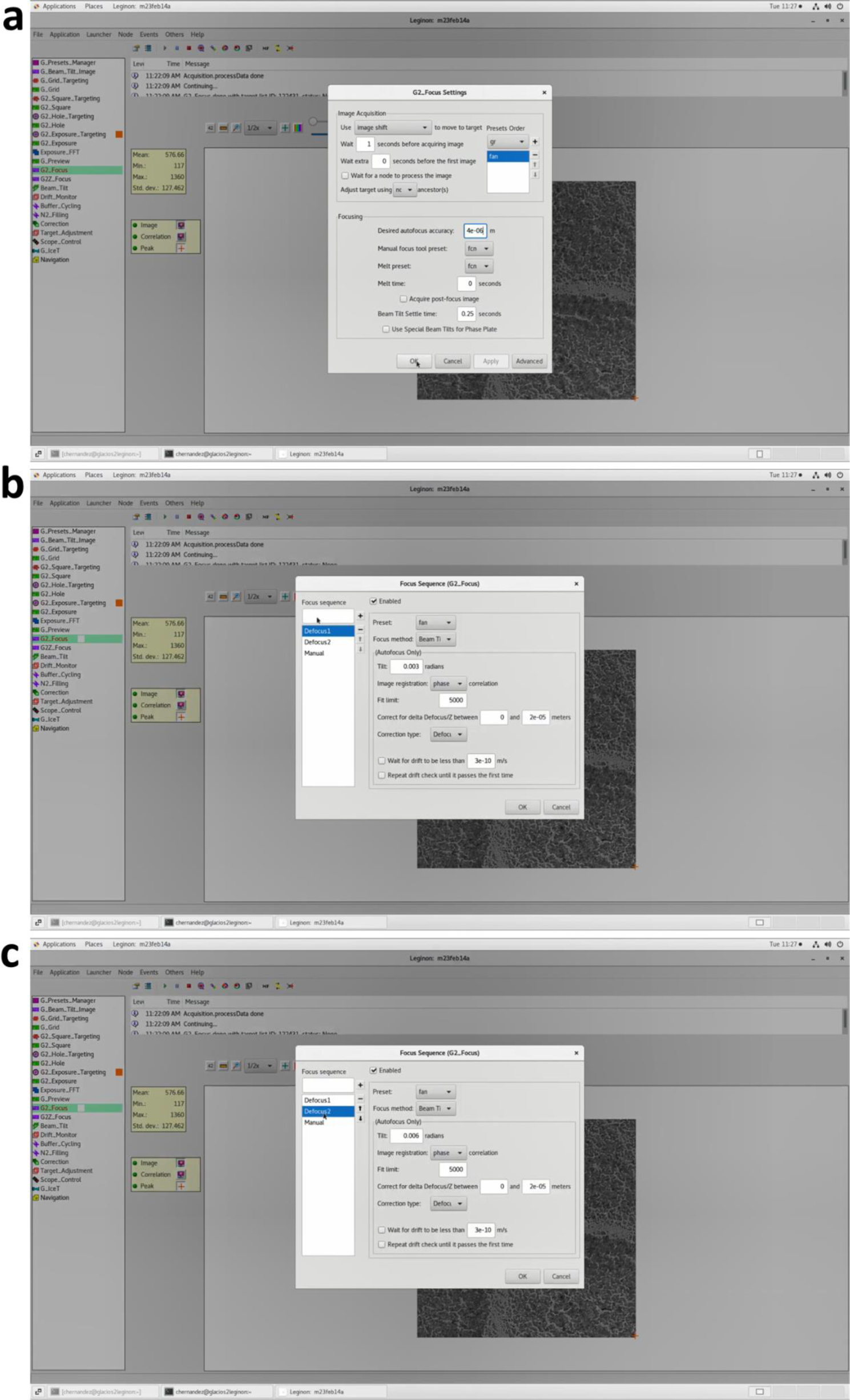
Focus settings and Focus Sequence settings for *Smart Leginon*.

**Supplementary Figure 5:**
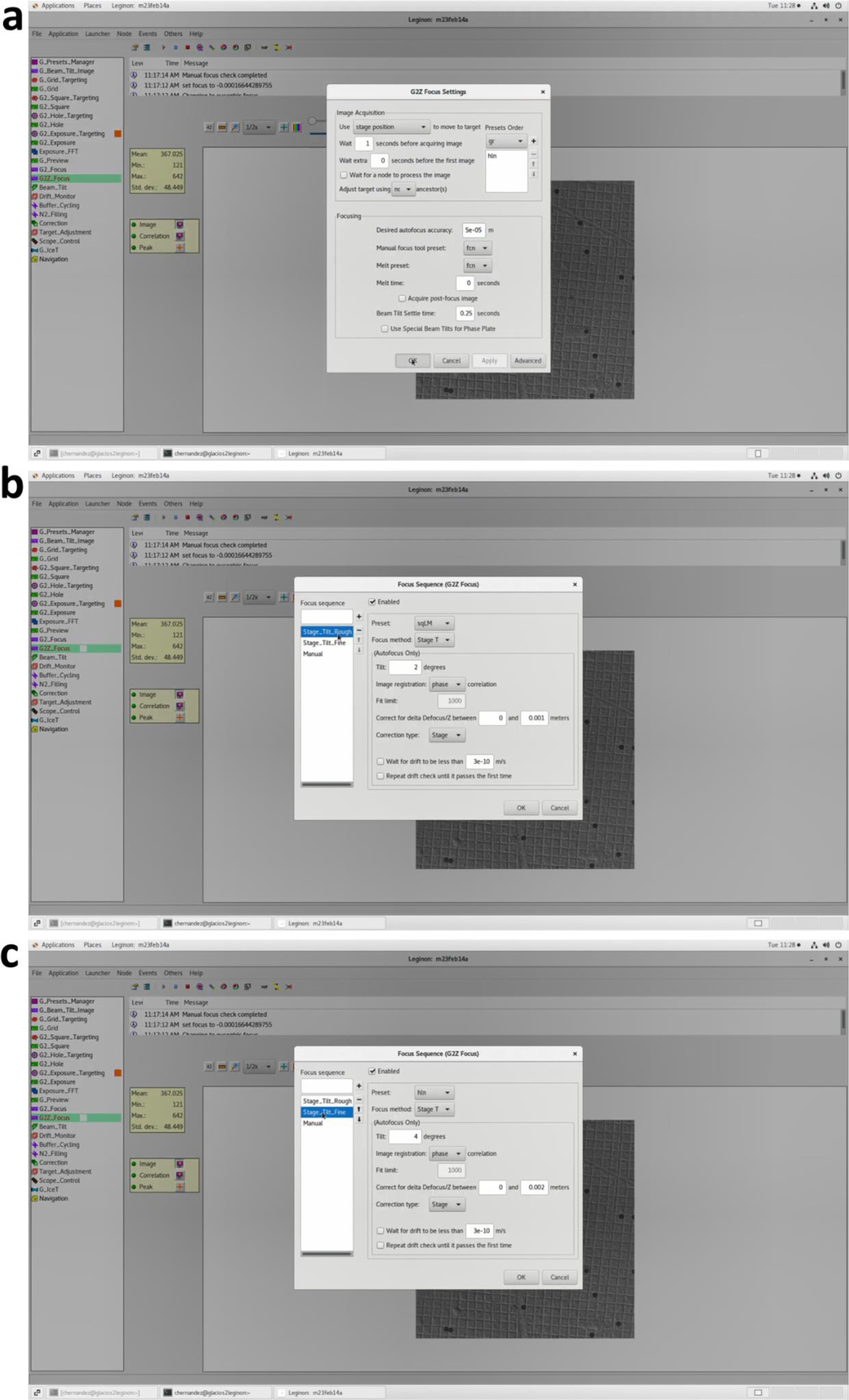
Z_Focus settings and Z_Focus Sequence settings for *Smart Leginon*.

**Supplementary Figure 6:**
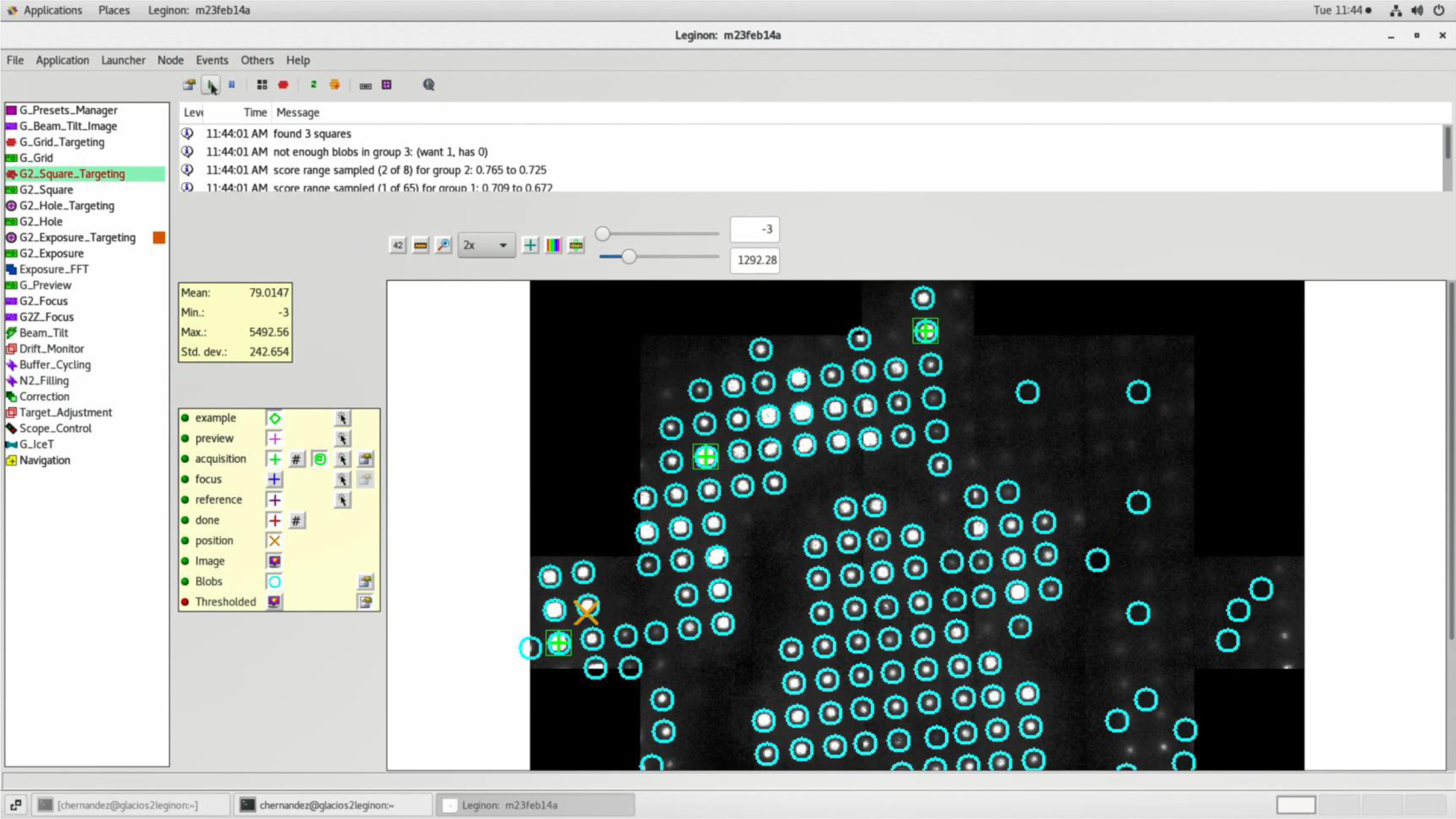
An example atlas after setting up *Smart Leginon* Square_Targeting parameters. Blue circles are blobs, green plus signs are acquisition locations, and the brown ‘x’ is the current stage location.

**Supplementary Figure 7:**
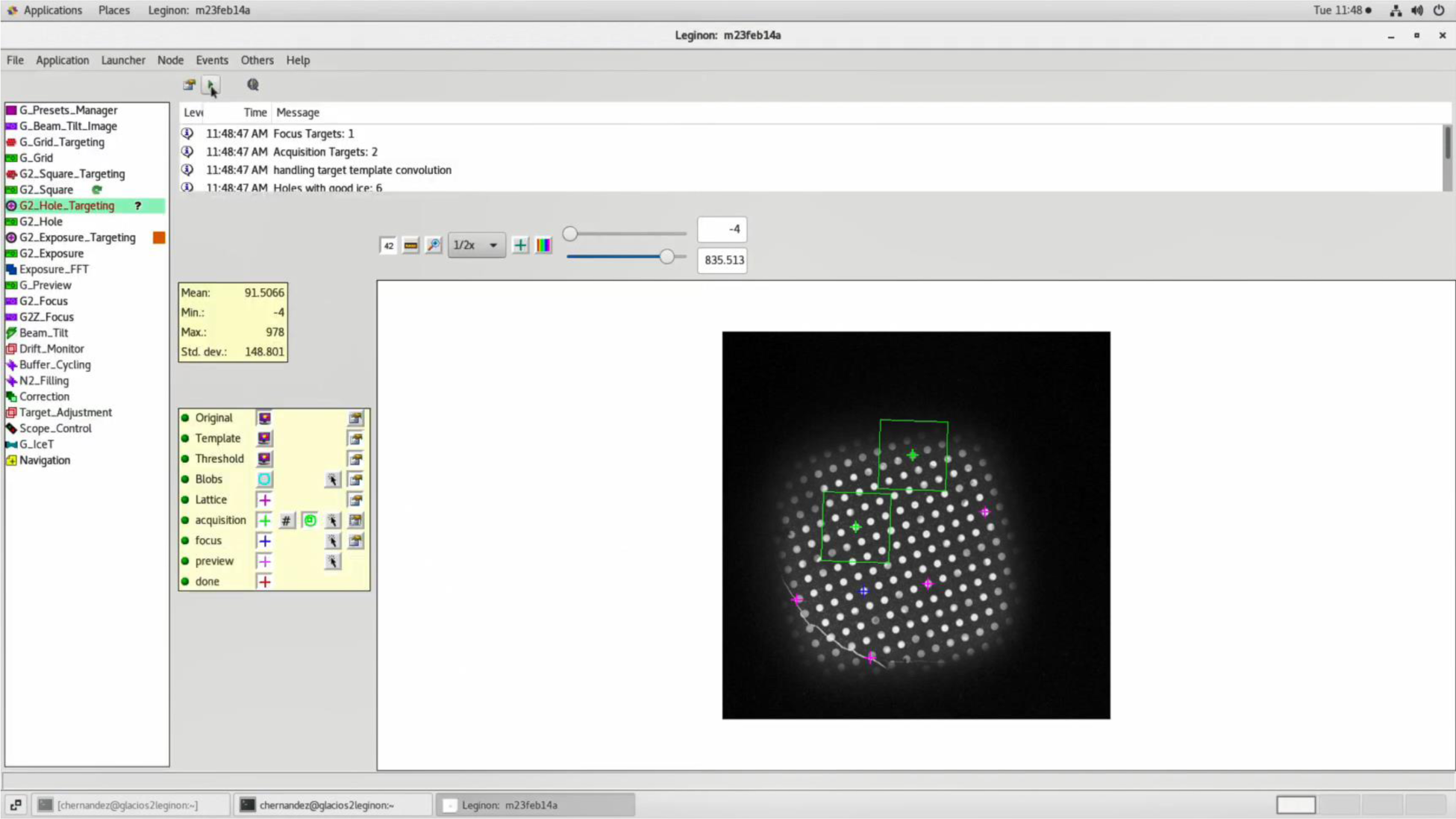
An example atlas after setting up *Smart Leginon* Hole_Targeting parameters. Purple plus signs are lattice locations, green plus signs with boxes are acquisition locations, and the blue plus sign is the focus location.

**Supplementary Figure 8:**
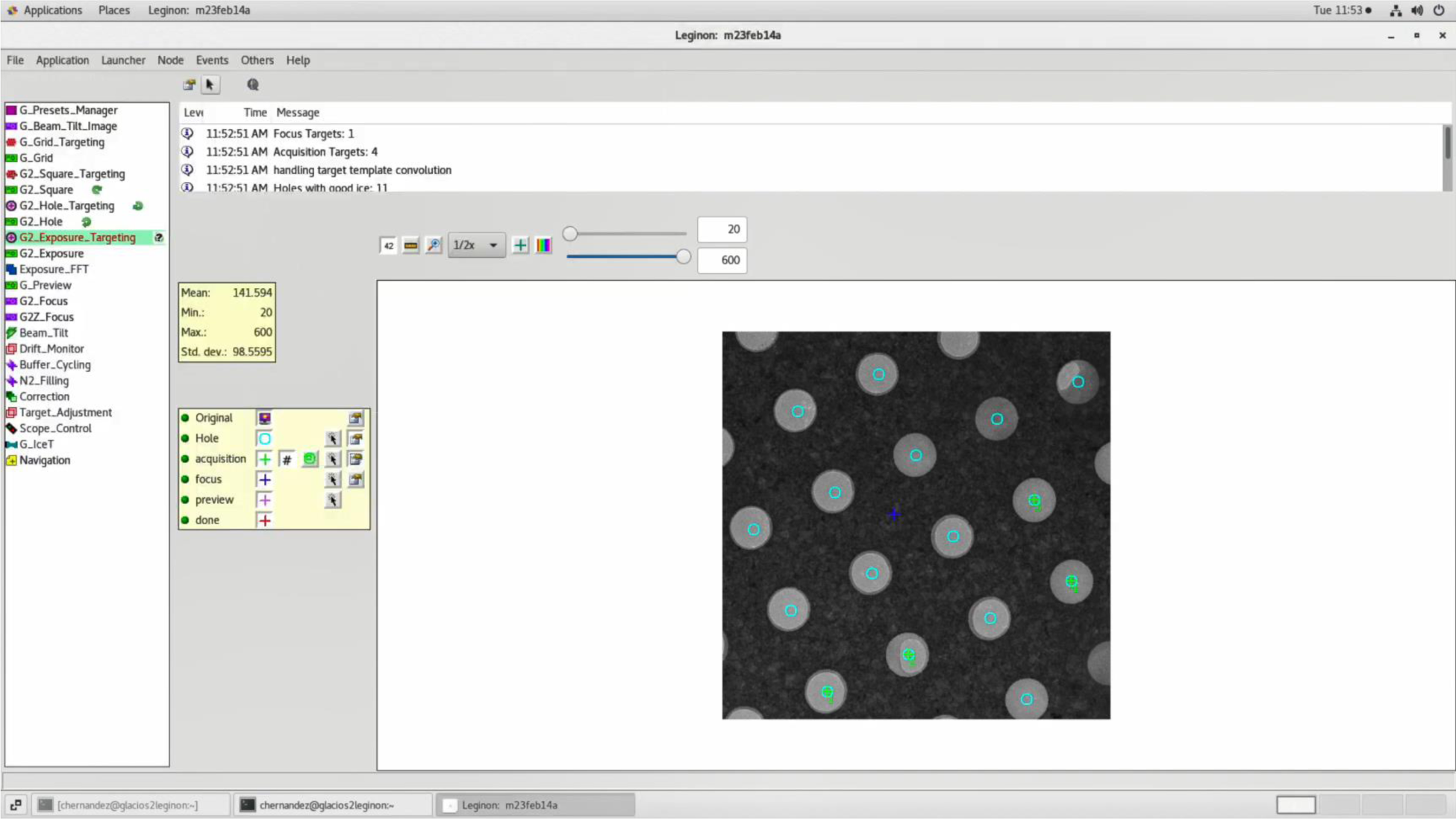
An example atlas after setting up *Smart Leginon* Exposure_Targeting parameters. Blue circles are blobs, green plus signs are acquisition locations, and the blue plus sign is the focus location.

**Table of Materials:**
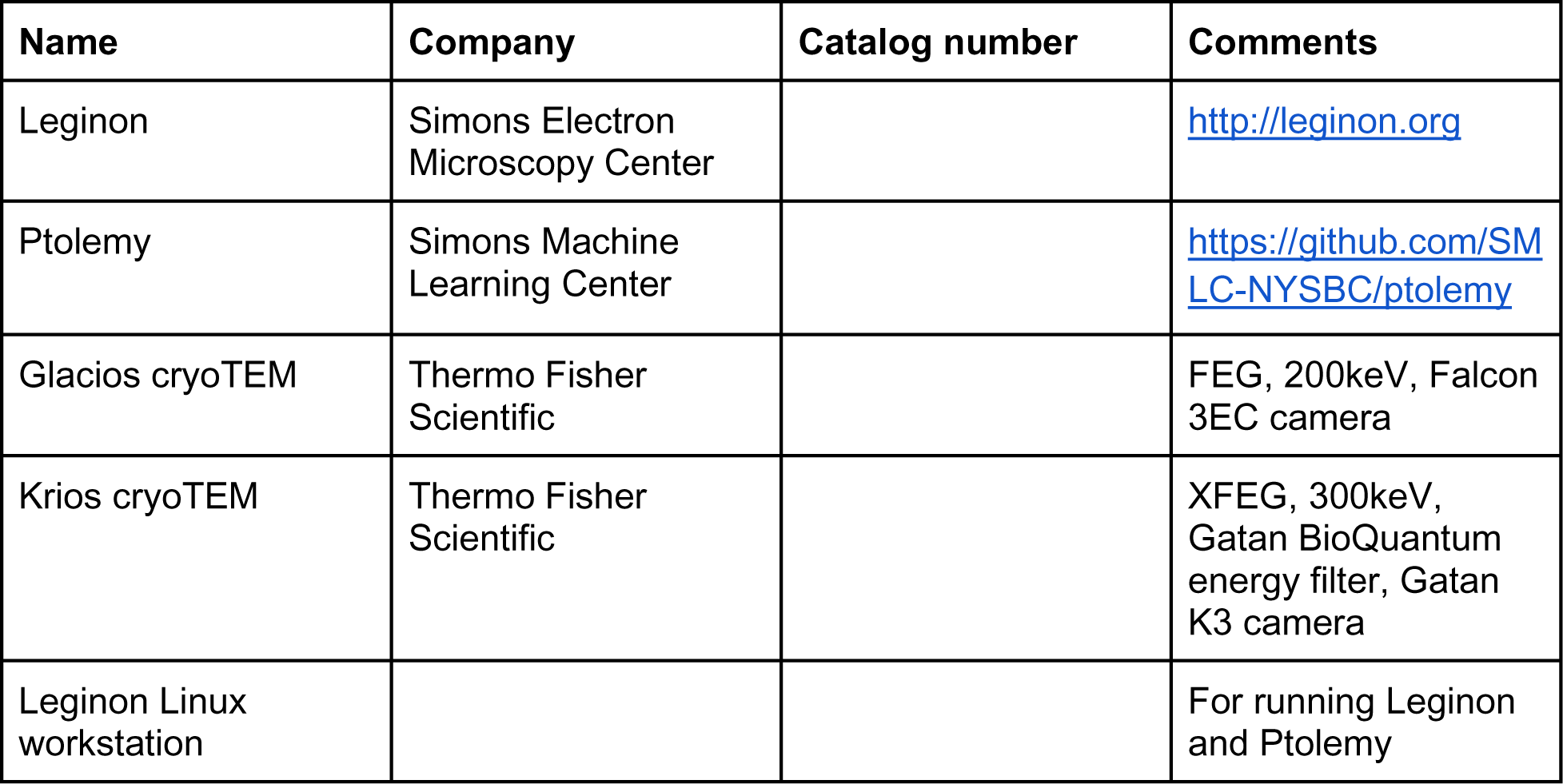
Required software and equipment for the protocol.

